# Use of the Elston-Stewart algorithm for the efficient calculation of exact pedigree-based Y-STR match probabilities

**DOI:** 10.64898/2026.08.14.744965

**Authors:** Janne Berger, Michael Krawczak, Dion Zandstra, Manfred Kayser, Arwin Ralf, Eva Scheurer, Amke Caliebe, Iris Schulz

## Abstract

The formal assessment of a genetic match between a suspect and some biological trace material is one of the key tasks of forensic genetics, particularly in cases of sexual offence. The analysis of Y-chromosomal short tandem repeats (Y-STRs) has proven especially useful in this context. For a long time, however, calculating the probability of a perfect Y-STR profile match under the defense hypothesis that the suspect was not the trace donor posed a great challenge. This was due to the inherent uncertainty about the population of alternative donors, the so-called ‘suspect population’. We recently proposed to resolve this controversy by systematically favoring the suspect and considering his close male relatives as the suspect population. However, since the mathematical framework developed for this purpose was simulation-based, its practical application turned out increasingly difficult with increasing pedigree size. Here, we present an adaptation of the so-called ‘Elston-Stewart algorithm’, originally developed for the linkage analysis of human genetic diseases, to allow calculation of exact match probabilities in a time that scales linearly with pedigree size. The adapted algorithm was implemented in a publicly available software tool, and its correctness was verified by the comparison of its output with the correct, analytical results obtained for selected example pedigrees. The new implementation mostly outperforms the simulation-based solution, albeit with the important exception of Y-STRs present in multiple copies. Given the increasingly prominent role of such ‘multicopy markers’ in forensic genetics, the complementary use of both approaches appears the most sensible strategy for the time being.

**Summary:** When the suspect’s DNA in a criminal case matches the DNA of a biological trace, the defense rightly wants to know the probability of this match, given that their client was not the trace donor. Calculating this probability has proven difficult in the past, particularly for male-specific genetic markers important to investigate sexual offenses. We previously suggested a simulation-based approach to resolve this problem, but the method’s performance decreased notably with increasing pedigree size. Here, we report the task-specific adaptation of an algorithm originally developed for medical genetics research that mostly resolves the runtime limitations of the simulation approach.

## Introduction

In many forensic cases, the DNA of a suspect needs to be compared to genetic material left at a crime scene to clarify whether the suspect contributed their DNA to the trace. Short tandem repeat (STR) markers, defined by the allelic number of repetitions of a specific short nucleotide motif, are routinely used for this purpose. While forensic DNA analysis usually focuses upon autosomal STRs, the informativeness of the latter may nevertheless sometimes be limited. More specifically, if the trace contains a mixture of few male cells and a large number of female cells, as is the case in many investigations of sexual offences, the male autosomal STR profile can be masked. In such instances, Y-chromosomal STRs (Y-STRs) can be used to identify potential male contributors to the trace without interference from the female DNA. Indeed, over the past 30 years, Y-STRs have become an indispensable tool, a “workhorse”, for this purpose (1, 2).

Handling DNA mixtures with multiple contributors in forensic casework is extremely challenging (3). For the sake of simplicity, we therefore assume in the following that a single origin of the Y-chromosomal DNA contained in a trace has been established beyond doubt. Under this condition, a suspect can usually be reliably excluded from donorship as long as his Y-STR profile differs from that of the trace at one or more of the markers included in the profile. However, if the two profiles match, i.e., if both samples exhibit the same allele at all markers, the probative value of that observation needs to be quantified in the form of the so-called ‘match probability’. As the name implies, this numerical value is intended to correspond to the probability of observing the match in question if the male component of the trace DNA originated from a person other than the suspect, selected at random from a population of plausible alternative donors.

To date, forensic genetic experts have routinely equated the match probability with the estimated frequency of the Y-STR profile of interest in an actual or hypothetical human population (4). Large national and international population reference databases, particularly the Y-chromosome STR Haplotype Reference Database YHRD (5), served as sources of the population genetic information required to this end. However, all these attempts suffered from an inherent uncertainty about the appropriate choice of said suspect population (6, 7). Whatever criteria are being used to define the latter, they may be challenged in court on the grounds that they are too unspecific for, and thus disadvantageous to, the suspect.

This dilemma prompted some of us to develop a novel mathematical framework for calculating match probabilities from the suspect’s male pedigree. Instead of viewing alternative trace donors as anonymous members of a vaguely defined suspect population, they were characterized by whether or not they were close male relatives of the suspect (8, 9). This approach was motivated by the fact that the increasing use of rapidly mutating (RM) Y-STRs, which are capable of distinguishing between father and son in almost 50% of cases (10, 11), makes exact profile matches between distantly related or unrelated males increasingly unlikely. Knowledge of the match probabilities for all untyped, potentially relevant family members should therefore both benefit police investigations and enable a fair and effective assessment of circumstantial evidence in court.

To promote its practical application, we recently implemented an extended version of the mathematical framework in a publicly available software tool named ‘MatchY’ (9, 12). As also originally described by Caliebe et al. (8), the current version of MatchY employs simulation-based importance sampling to calculate the required match probabilities, i.e., the probabilities of specific assignments of the suspect Y-STR profile to untyped pedigree members, taking into account all relevant blood relationships. The importance sampling method was chosen for this purpose because including more distant relatives of the suspect in the analysis also would have increased the number of common ancestors to be considered so that the deterministic calculation of exact match probabilities was deemed unrealistic given realistic pedigrees sizes. However, as is to be expected with any simulation-based approach, the speed at which MatchY calculations converge to the true value decreases in scenarios involving multiple markers, greater profile diversity, and a greater number as well as a less favorable positioning of the untyped pedigree members (12).

These circumstances originally threatened to limit the practical usability of the MatchY tool. However, during the further development of the underlying algorithm, we found that reversing all father-son relationships connecting the suspect to the most recent common ancestor (MRCA) of the pedigree members significantly increased simulation efficiency and, consequently, the speed of convergence of the MatchY calculations. Perhaps even more important, by turning the suspect into the MRCA and conditioning the calculations on his Y-STR profile, exact probability calculations became possible again.

In fact, an algorithm developed in 1971 by early genetic epidemiologists Robert C. Elston and John Steward (13) for human genetic linkage analysis offers a potential solution to the scaling problems of MatchY. The Elston-Stewart (or ES) algorithm, as it has since been known, enables the recursive calculation of pedigree-based probabilities in a way that scales exponentially with the number of markers, but only linearly with pedigree size. Furthermore, conditioning the calculations on the suspect’s known Y-STR profile implies that all inheritance patterns within the pedigree are determined exclusively by mutation events. Since mutations at different Y-STR markers can be assumed to occur independently (14, 15), the first characteristic of the ES algorithm (i.e., exponential scaling with marker number) becomes irrelevant in the present setting: Marker-specific match probabilities can be calculated individually and subsequently multiplied to obtain the overall match probability. The second characteristic (i.e., linear scaling with pedigree size), on the other hand, is an obvious key to solving the looming efficiency dilemma of MatchY. Therefore, we describe below the implementation of a simplified version of the ES algorithm, specifically tailored to the requirements of the calculation of exact pedigree-based Y-STR match probabilities.

## Material and Methods

### Background and definitions

The primary purpose of this and previous work (8) has been to calculate the match probability between the Y-STR profile of a suspect and that of given untyped members of his male pedigree. Note that the mean of these probabilities equals the match probability of a randomly drawn untyped member, which is conceptually equivalent to the classic population-based match probability.

Each member-specific match probability is calculated from the probability of the Y-STR profiles observed within the pedigree (or the ‘pedigree probability’, for short), taken once in its original form and once with the profile of the untyped member of interest set equal to the suspect profile. By definition, the ratio of these two figures equals the sought-after match probability P(i) of the i^th^ untyped male, i.e.

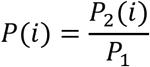

Here, P_1_ is the original pedigree probability and P_2_(i) is the pedigree probability with the suspect profile assigned to the i^th^ untyped male.

From now on, it suffices to consider just a single Y-STR. This is because the pedigree probability is always calculated conditional on a known founder haplotype so that the overall pedigree probability corresponds to the product of the single-marker pedigree probabilities (see Introduction). For the sake of simplicity, we also focus on a single-copy Y-STR with integer-valued alleles bounded by values minall and maxall, respectively. We denote by a(i)ɛ{0,minall,..,maxall} the Y-STR allele of the i^th^ (typed or untyped) pedigree member, where a(i)=0 means that the i^th^ member is not typed. In addition, p(j→k) denotes the probability that paternal allele j is inherited by a son as allele k. Note that the current implementation of the ES algorithm assumes a symmetric one-step mutation model with mutation rate µ, i.e., that p(j→k)=µ/2 for k=j±1, p(j→k)=1-µ for k=j, and p(j→k)=0 otherwise.

### The ES algorithm adapted

Let β(i,j) denote the conditional probability of the Y-STR profiles of all descendants of the i^th^ male, given that the i^th^ male carries allele j. We first consider a two-generation pedigree with a father (0^th^ male) and his n sons (1^st^ male to n^th^ male). Then

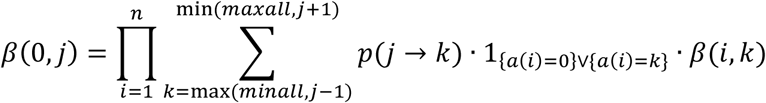

If the i^th^ male (i>0) has no descendants, β(i,k) is set equal to unity.

In the general case, a pedigree comprises m+1 generations (numbered 0 to m), with a total of n+1 males (numbered 0,..,n) and n(j) males belonging to the j^th^ generation. The (typed) MRCA is designated the 0^th^ male and, by definition, is the sole member of the 0^th^ generation. Then our goal is to calculate the overall pedigree probability β(0,a(0)).

Without loss of generality, we can assume that the numbering of pedigree members is by generation and by sibship within generations. This means that, (1) if the i^th^ male belongs to an earlier generation than the j^th^ male, then i<j, and (2), if the i^th^ and j^th^ male (i<j) belong to the same sibship, then any k^th^ male with i<k<j also belongs to that sibship. Let f(j) be the smallest index of males belonging to the j^th^ generation, which means that the indexes of the j^th^ generation lie between f(j) and f(j)+n(j)-1=f(j+1)-1. Let ns(i) be the number of sons of the i^th^ male, and let id(i,j) be the index of the j^th^ son of the i^th^ male.

Following the principle of the ES algorithm, the pedigree probability β(0,a(0)) is calculated recursively from the bottom to the top of the pedigree as follows:

for j_1_=f(m) to n do for j_2_= minall to maxall do β(j_1_,j_2_):=1 for j_1_=m-1 to 0 do for j_2_=f(j_1_) to f(j_1_)+n(j_1_)-1 do if a(j_2_)=0 then for j_3_=minall to maxall do

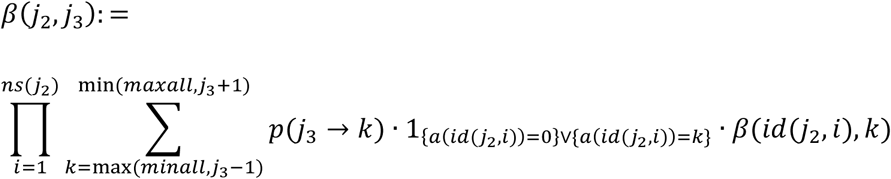

else

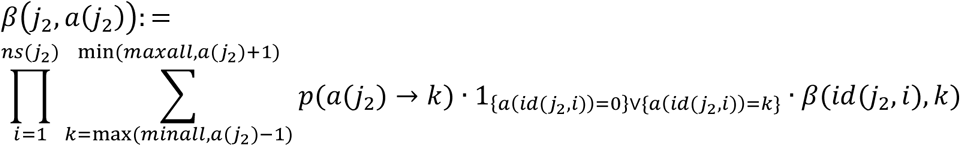

### Algorithm implementation

The adaptation of the ES algorithm described above has been implemented in Python 3.14.3 (16) using the Python standard library. Pedigree and genetic information are provided by the user as csv files with columns *male_id, generation, father_id*, and one *allele_<LOCUS>* column per marker. A command-line interface (cli.py) allows direct application of the algorithm to correctly specified pedigrees.

Central to the implementation is the *compute_match_probs* function, which calculates the required pedigree probabilities by two unidirectional passes over the pedigree that avoid redundant calculations otherwise caused by untyped individuals (17). A first (‘inside’) pass, starting at the terminal leaves of the pedigree and moving upwards to the MRCA (i.e., the suspect), calculates the original pedigree probability, P_1_, by so-called ‘recursive backward induction’. A second (‘outside’) pass, starting at the MRCA and moving downwards to the terminal leaves, complements the inside pass and yields the individual-specific pedigree probabilities, P_2_(·), directly without a need to repeat the backward induction for each untyped pedigree member (17). The *compute_match_probs* function also calculates the mean of the P_2_(·) values, which corresponds to the match probability of a randomly drawn, untyped pedigree member.

The source code of the ES algorithm implementation is publicly available under the MIT license at https://github.com/JanneBerger/ExactYSTR.

### Implementation correctness

To verify the correctness of the ES algorithm implementation, Y-STR match probabilities were computed for five example pedigrees with different allele assignments (Figure 1), adapted from our original mathematical framework report (8). The results were then compared to both the analytical (and hence correct) match probabilities and the simulation-based results of MatchY.

**Figure 1:**
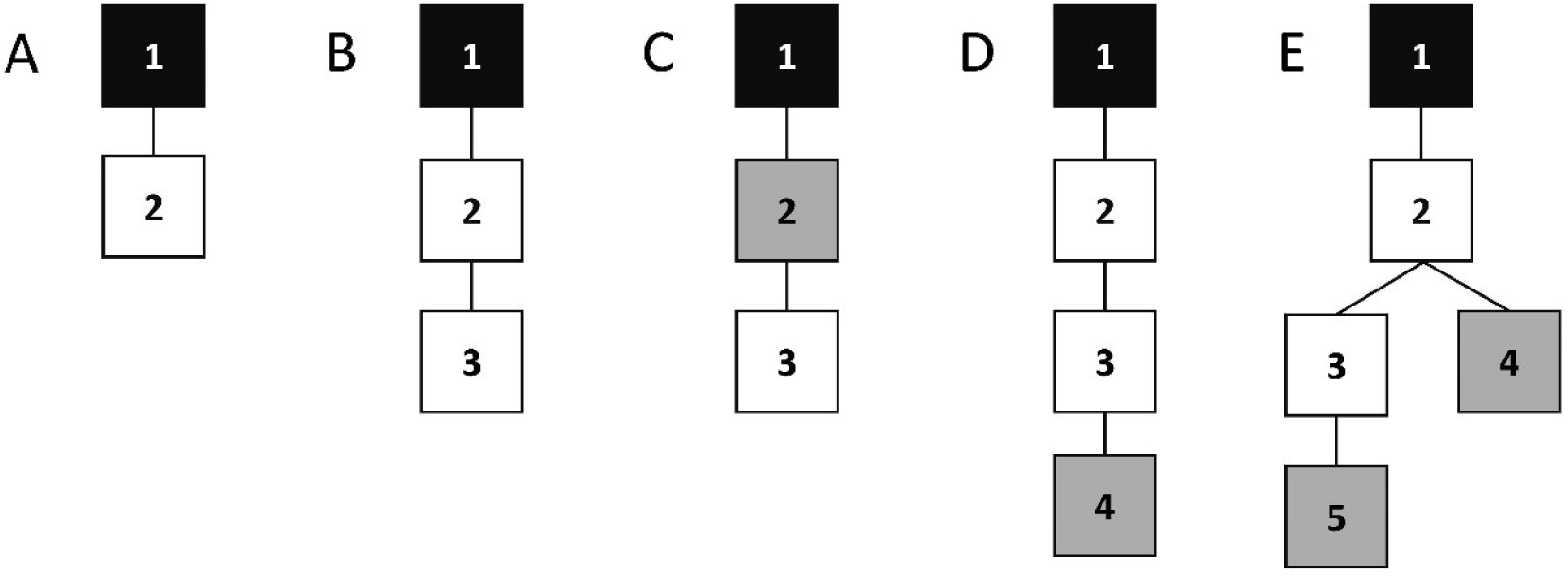
Example pedigrees used to verify the correctness of the ES algorithm implementation (adapted from (8)). Black square: most recent common ancestor (MRCA); grey squares: typed males; white squares: untyped males. Due to the routine restructuring of the pedigree in MatchY, the MRCA corresponds to the suspect and is therefore always typed. Alt text: Example male pedigrees, featuring a typed suspect and some of his close relatives, that were used to verify the correctness of the ES algorithm implementation.

### Computational efficiency

The computational efficiency of the ES algorithm adaptation was evaluated by a Python script (*runtime_analysis.py*) that analyzed the runtime required under various scenarios in terms of pedigree size and complexity as well as marker number. The runtime was measured with *time.perf_counter()* as the mean outcome of 20 *compute_match_probs* calls with different random selections of untyped pedigree members (for details, see below). An additional warm-up call was performed prior to these measurements to ensure that initialization effects related to Python execution and CPU caching did not affect the actual runtime measurement.

To evaluate computational efficiency, the following parameters were considered: the pedigree size (i.e., the number of pedigree members) n_tot_, the number of untyped males n_unt_, and the number of marker alleles n_all_. In addition, it was verified whether the runtime increases linearly with the number of loci n_loc_ as expected. Each parameter was varied individually (Table 1) while the other parameters were held constant at typical values. All measurements were repeated for three pedigree topologies, namely a linear chain (number of offspring per male n_off_=1), a ternary tree (n_off_=3), and a quinary tree (n_off_=5), to investigate how the runtime depends on the structure of the pedigree, in addition to its size. A linear model was fitted to the measured runtimes using *numpy.polyfit* to assess whether or not the runtime increased linearly with the respective parameter.

**Table 1:** Parameter sweeps used for the evaluation of computational efficiency.

| Varied parameter | Fixed parameters |  |  |  | Parameter sweep |
| --- | --- | --- | --- | --- | --- |
| | $n_{\text{tot}}$ | $n_{\text{unt}}$ | $n_{\text{all}}$ | $n_{\text{loc}}$ | |
| $n_{\text{tot}}$ | -.- | 30 | 28 | 1 | 50, 75, 100, 125, 150, 175, 200, 250, 300, 350, 400, 450, 500 |
| $n_{\text{unt}}$ | 200 | -.- | 28 | 1 | 1, 2, 5, 10, 20, 30, 50, 75, 100, 125, 150, 175, 199 |
| $n_{\text{all}}$ | 200 | 50 | -.- | 1 | 4, 6, 8, 10, 14, 18, 22, 26, 28 |
| $n_{\text{loc}}^*$ | 50 | 25 | 28 | -.- | 1, 2, 3, 5, 8, 10, 15, 20 |

\*Since *compute\_match\_probs* is called per locus, runtimes were summed over all $n_{\text{loc}}$ loci for efficiency assessment.

For each combination of topology and parameter setting, pedigrees were generated with Python’s built-in *random.sample* function, randomly selecting n_unt_ males from the non-root members of the pedigree and designating them as ‘untyped’. Every pedigree construction was repeated five times with the same random seed, and the median runtime measured was used for further analysis in order to reduce technical noise. Calculations were performed on a laptop with an Intel Core i9-13900H processor, 64 GB RAM, and running Windows 11 Pro.

Since, in this way, all non-root males had an equal chance of being selected, the untyped members were naturally distributed evenly across all generations. While we were aware that this set-up may be highly unrealistic, particularly if a large number of non-selected (i.e., typed) males belong to early generations, it was both easy to implement and systematic, and hence considered appropriate for the comparative evaluation of computational efficiency.

## Results

### Implementation correctness

To verify the correctness of the ES algorithm implementation, the computational results obtained for five exemplary pedigrees (Figure 1), using different allele assignments to typed males in some cases, were compared with both the analytical (i.e., correct) results and the simulation-based output of MatchY (Table 2). For all pedigrees and allele assignments, the implementation of the ES algorithm yielded exactly the analytical match probabilities, thereby confirming the correctness of the implementation. The MatchY results were close to the exact values but not identical to them, as was to be expected for these simulation-based calculations.

**Table 2:**
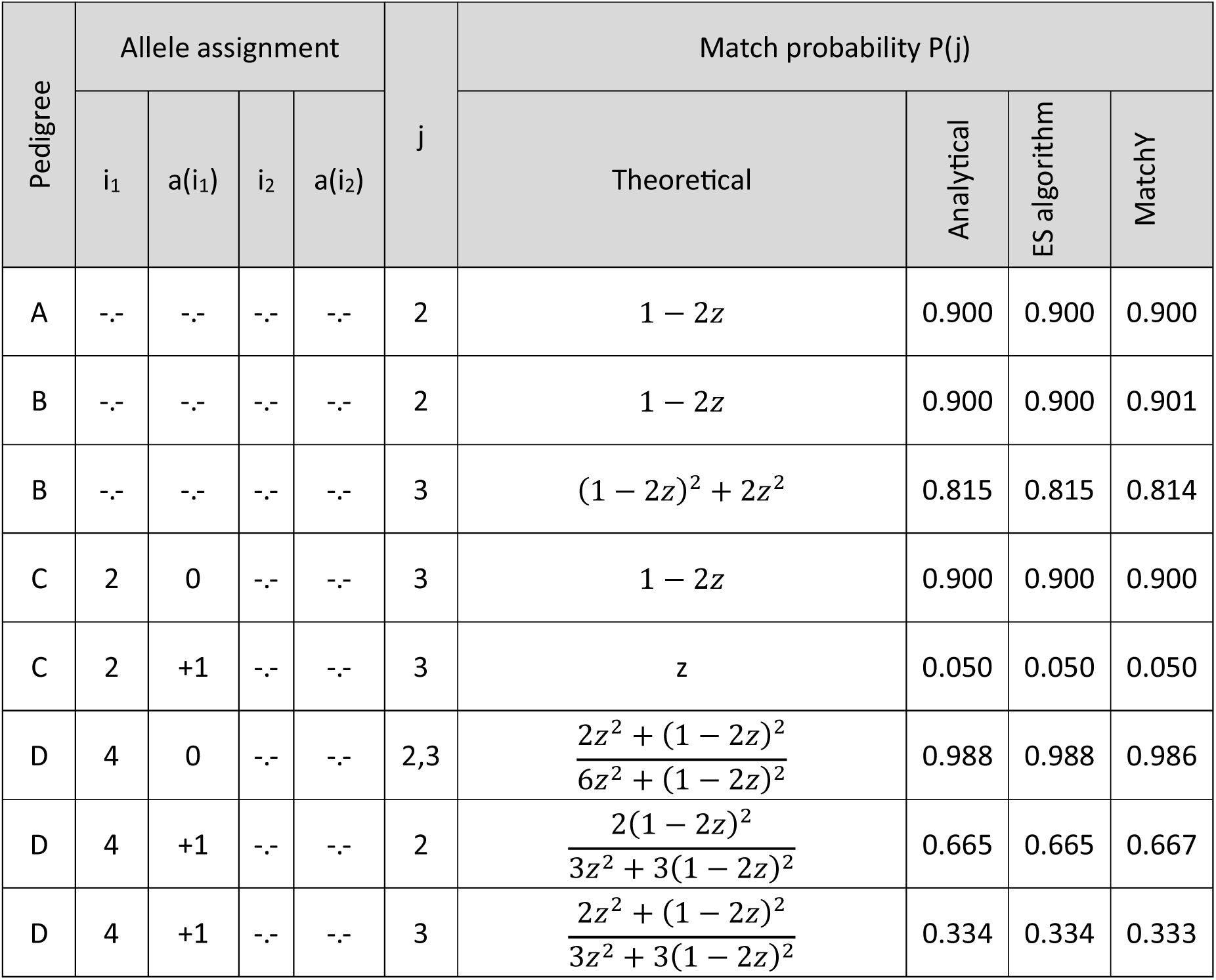
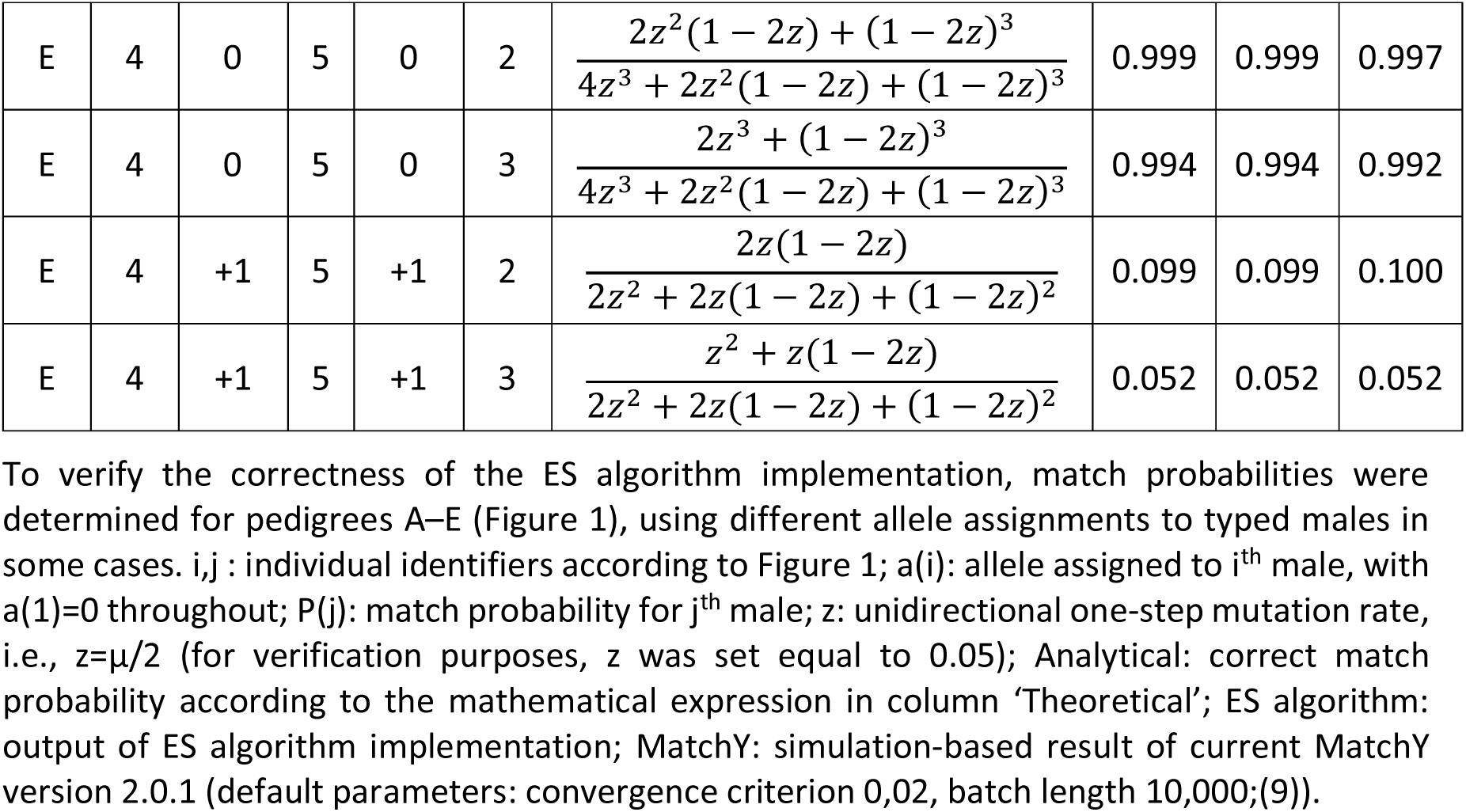
Verification of correctness of ES algorithm implementation.

### Complexity and runtime

The effects of various parameters (Table 1) on the runtime requirements of the ES algorithm implementation were investigated utilizing three different pedigree topologies, differing by their degree of branching. Specifically, a male had either exactly one son (linear chain), exactly three sons (wide pedigree) or exactly five sons (very wide pedigree). For all four parameters, the required runtime increased more or less linearly with the respective parameter (Figure 2). Linear chain pedigrees consistently required the longest runtime with approximately 30 ms at n_tot_=500 (Fig. 2a) and approximately 14 ms at n_unt_=199 (Fig. 2b). The calculation of pedigree probabilities for wider topologies was consistently found to be faster.

**Figure 2:**
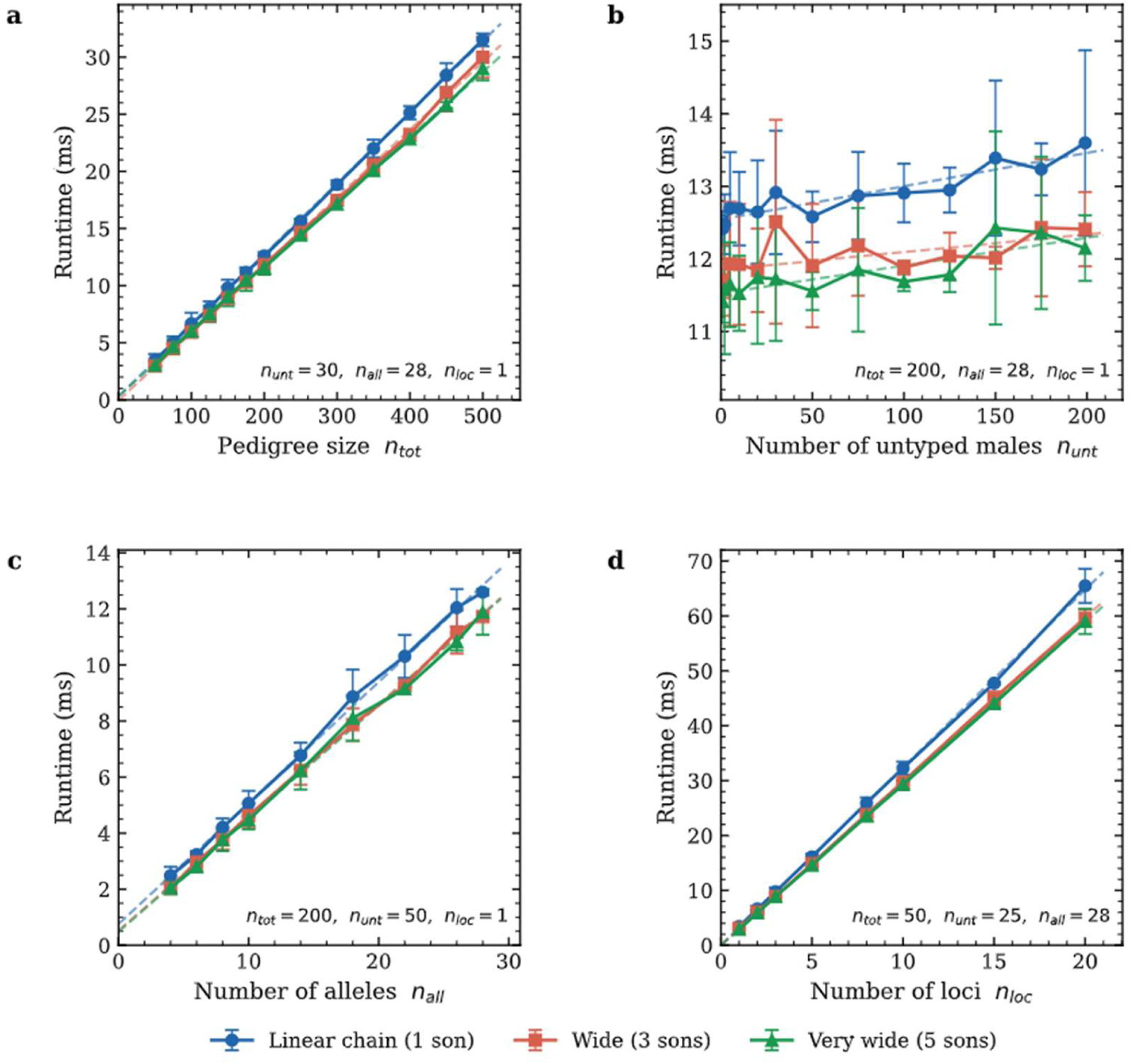
Runtime requirements of the ES algorithm implementation. Runtime was analyzed as a function of four different parameters, namely the pedigree size n_tot_ (a), the number of untyped males n_unt_ (b), the number of alleles n_all_ (c), and number of loci n_loc_ (d). Three different pedigree topologies were considered: a linear chain (blue, 1 son per male), a ternary tree (red, 3 sons per male), and a quinary tree (green, 5 sons per male). Each data point represents the mean runtime from 20 calls of routine *compute_match_probs* with the same parameter setting and pedigree topology, but different random selections of untyped members. For each call, the median of five identical replicates with the same random seed was used to account for potential measurement noise. Fixed parameter settings are given in an inset within each graph. Dashed lines indicate linear fits obtained using the least-squares method. Alt text: Relationship between the runtime of the ES algorithm and four parameters: pedigree size, number of untyped males, number of alleles, and number of typed markers. Each relationship is presented for three pedigree topologies, namely a linear tree, a ternary tree, and a quinary tree, supplemented by a fitted linear model.

Across all parameter combinations investigated, the runtime did not exceed 70 ms, indicating that the practical application of the ES algorithm is computationally feasible, even with large pedigree sizes and extensive marker sets.

## Discussion

“The right way is not always the popular and easy way.”

Margaret Chase Smith (US-American politician, 1897-1995).

Since the introduction of Y-STRs in forensic practice more than 30 years ago (1, 2), the formal assessment of matches between the marker profiles of a suspect and a trace has followed an intuitive, and therefore widely accepted, paradigm: If the suspect was not the trace donor, then the donor must have been an (unknown) member of a population of equally probable alternative donors - the so-called ‘suspect population’. As plausible as this perspective may be, it has a crucial drawback. Since suspect populations can be defined in various ways, a criterion is required for this purpose to which all parties involved can agree, including the suspect himself and his lawyers. This has consistently proven to be a difficult task because the match probability is naturally higher, and thus more advantageous for the suspect, the greater the genetic similarity between the suspect and other members of the suspect population. Consequently, none of the currently used, primarily geography-based and relatives-excluded criteria are immune to the defense criticism that they may disadvantage the suspect.

A criterion was thus required that is largely immune to these objections. The increasing use of rapidly mutating (RM) Y-STR markers in forensic practice proved timely in this regard as it allowed the focus to be placed on distinguishing between the suspect and his close male relatives. This option would indeed be very helpful because, regardless of how alternative suspect populations are defined, none of them would be genetically as close to a given male suspect in terms of his Y-chromosomal genetics as his sons, brothers, and patrilineal cousins. Moreover, kinship ties are typically well-documented in modern times, providing the investigative authorities with a better basis for their investigations than geographical characterizations such as “Northern European” or “resident within a 50-km radius”.

The above considerations led some of us to undertake a theoretical treatment of pedigree-based Y-STR match probabilities (8) and to subsequently implement the resulting mathematical framework in the software tool MatchY (9, 12). Both the framework and the tool drew upon a simulation method known as ‘importance sampling’, which is frequently employed for complex probability calculations. However, while importance sampling worked in principle for our purposes as well, it soon became apparent that its reliance on simulations could potentially limit the practical usability of MatchY. Consequently, as we further developed both the framework and the tool, we sought a way to calculate pedigree-based match probabilities directly, thereby avoiding excessive runtimes at least for large pedigrees and high levels of Y-STR profile diversity.

Fortunately, the problem in question already existed half a century ago, albeit in a different context, when the first linkage analyses of monogenic disorders required a probabilistic assessment of the co-segregation of disease states and marker genotypes in extended pedigrees. The most prominent answer to this challenge was the Elston-Stewart (ES) algorithm, first published in 1971 (13). The fundamental principle of the ES algorithm was to successively integrate the genetic and phenotypic information about relevant offspring into their respective parents, proceeding backwards through generations to the root of the pedigree. In its general form, the algorithm is far more complex than required for the analysis of Y-chromosomal markers, where generations are clear-defined and non-overlapping, and where no kinship loops can occur. Nevertheless, the ES algorithm offered an ideal approach for the calculation of exact pedigree-based Y-STR match probabilities within a practical timeframe, which is why we adapted and implemented it for this purpose.

The efficiency of the ES algorithm derives from the principle of ‘dynamic programming’ (DP), a fundamental concept in computer science first introduced in the 1950s (18). The core idea of DP is to decompose a complex problem into overlapping subproblems, to solve each subproblem once, and to reuse the results so as to avoid redundant re-computation. Elston and Stewart (13) recognized that pedigree likelihood calculations have exactly this structure: the conditional genotype probability of a descendant of a given pedigree member does not depend upon the rest of the pedigree, which means that it has to be computed only once and that, consequently, the runtime scales only linearly with the pedigree size.

Here, we not only confirmed this DP-derived property for our implementation of the ES algorithm but also demonstrated linear runtime scaling with respect to other performance-relevant parameters, specifically the number of untyped pedigree members, the number of markers, and the number of alleles per marker. The algorithm should therefore outperform the simulation-based routines of MatchY with growing pedigree size and increasing marker number. However, there are two important exceptions that may still make simulation with MatchY indispensable, at least for the time being.

First, the current implementation of the ES algorithm is limited to single-copy Y-STRs, and an extension to multi-copy markers, including many RM Y-STRs, seems difficult to achieve efficiently. As long as alleles cannot be unambiguously assigned to individual marker copies, genotypes can only be coded as unordered lists between which mutations are no longer symmetric. For example, even if both copies of a Y-STR have the same symmetric mutation rate, transition {10,10}→{10,11} is twice as probable at the genotype level as {10,11}→{10,10}. This is because the former transition can result from either of the two possible 10→11 mutations whereas the latter requires that (single) allele 11 mutates to 10. Extending the ES algorithm to multi-copy markers would therefore entail tracking all father-son inversions, introduced during the initial restructuring of the pedigree, through all subsequent calculations. Furthermore, a significantly more complex genotype indexing scheme would be required to define mutation probabilities and to account for untyped pedigree members. In practice, it may therefore be advisable to initially limit the calculation of match probabilities to the available single-copy markers, using the ES algorithm implementation. If the results obtained in this way do not meet the case-specific requirements, the simulation-based method implemented in MatchY should be applied to additional multi-copy markers, and the results combined.

Second, the mathematical framework underlying MatchY was originally focused on the probability of a specific number of matches within the pedigree. Whether and to what extent these numbers are relevant in practice may be an open question. However, calculating them using the ES algorithm would be difficult to implement because the process cannot be carried out through the sequential consideration of untyped pedigree members. Instead, every possible combination of potentially matching males would have to be considered, the number of which grows exponentially with the number of untyped males. Therefore, if the probability of a specific number of matches is a priority, it would have to be determined either through computationally intensive, repeated executions of the ES algorithm, or by utilizing the simulation-based functionality of MatchY. As was mentioned above, however, the classical population-based match probability also merely informs the court of the chance that a single, randomly selected member of the suspect population carries the suspect’s haplotype. The pedigree-based equivalent of this quantity is the mean match probability across all untyped pedigree members, which the current implementation of the ES algorithm can readily provide.

To benefit from the forensic use of pedigree-based Y-STR match probabilities, it is not strictly necessary to type all, or even a larger number of, the male relatives of the suspect. At present, doing this would likely entail considerable logistical challenges and, particularly in the families involved, may meet with significant resistance. Furthermore, in most legal systems, it would first be necessary to determine whether extensive genetic testing of presumably innocent individuals is permissible at all. Nevertheless, it is worth noting at this point that our approach may well prove advantageous even if only a small number of individuals other than the suspect were typed, especially if the family structure is reliably known and typing includes strategically important pedigree members. For example, two or more mismatches between the suspect and his father, as would be expected in approximately ≥10% of father-son pairs for RM Y-STRs (19), would effectively limit the possibility of a ‘perfect’ match to the descendants of the suspect.

Ultimately, it is to be hoped that the suggested divide of the suspect population into close relatives on the one hand, and distant relatives or non-relatives on the other, will overcome current obstacles and gain broad acceptance by the forensic genetic community in the near future. Fully in the spirit of the opening quote by Margaret Chase Smith, we are convinced that this is the right way for evaluating Y-STR profile matches, even if it currently appears neither popular nor easy.

## Acknowledgements

Generative AI (Claude Sonnet 5, Anthropic) was used for code development and code documentation. All code was reviewed, tested and validated by the authors.

